# No adaptive plasticity in the heat tolerance of lizard sperm

**DOI:** 10.64898/2026.07.21.739836

**Authors:** Wayne Wen-Yeu Wang, Benjamin Pethe, Abigail P. Ward, Waverly Wood, Colin T. Smith, Alanna J. Frick, Shannan S. Yates, Alex R. Gunderson

## Abstract

1. Adaptive phenotypic plasticity can help organisms cope with global warming by increasing their heat tolerance, yet most research has focused on whole-organism traits. Much less is known about thermal plasticity at the gamete level despite increasing evidence that reproduction, and particularly sperm function, is especially sensitive to heat.
2. We tested for plasticity in sperm heat tolerance in two ways: via male thermal acclimation (pre-ejaculate stage) and via sperm cell heat hardening (post-ejaculate stage) in brown anole lizards (*Anolis sagrei*). We also measured plasticity in several other sperm and ejaculate traits in response to male thermal acclimation.
3. To test for effects of male thermal acclimation, adult males were exposed to one of two ecologically realistic, fluctuating temperature regimes that either mimic cool spring conditions or projected future summer conditions for eight weeks. Throughout this period, we repeatedly measured sperm heat tolerance (LT50), baseline motility, and sperm count. We also measured sperm morphology at the end of thermal acclimation. We predicted that males in the warmer treatment would produce more heat tolerant sperm.
4. To test for post-ejaculate heat hardening, we compared the heat tolerance of ejaculated sperm that either did or did not experience a high but non-lethal temperature prior to heat tolerance measurement.
5. We found no plasticity in sperm heat tolerance due to either male thermal acclimation or sperm cell heat shock. Long-term thermal acclimation of males did not increase sperm heat tolerance, nor change motility, sperm count, or sperm morphology. We also found no evidence for post-ejaculate heat hardening, as exposing sperm cells to mild sublethal heat shock did not enhance sperm heat tolerance.
6. Our results indicate that gametic traits have limited capacity for plastic adjustment to thermal stress, which is broadly consistent with the low levels of thermal plasticity found for whole-organism thermal tolerance across ectotherms. This highlights the vulnerability of reproductive traits to rising temperatures and the importance of evolutionary and behavioral responses to buffer organisms from climate change.

## 1. Introduction

Under current projections, global temperatures are expected to rise 2 to 3 °C by the end of the century (IPCC, 2014). The increase in both mean global temperature and the frequency of extreme heat events caused by climate change is a major threat to biodiversity (Christidis et al., 2015; Meehl and Tebaldi, 2004; Sinervo et al., 2010; Pottier et al., 2025). Ectotherms, whose physiology is directly shaped by external conditions, are particularly vulnerable to these changing conditions, with many species predicted to face increased risk of overheating, range contractions, population declines, and extinctions (Murali et al., 2023; Urban, 2024; Sinervo et al., 2010; Pottier et al., 2025).

Adaptive thermal plasticity is one of the key mechanisms that could allow ectotherms to cope with rapid temperature change (Huey et al., 1999, 2012; Angilletta, 2009; Gunderson et al., 2017). Thermal plasticity refers to within-generation phenotypic responses to variation in temperature and is considered adaptive when it improves fitness (Angilletta, 2009; Wilson and Franklin, 2002; Ghalambor, 2007). In the context of thermal physiology, adaptive plastic responses to high temperatures involve increased tolerance for heat (Angilletta, 2009; Gunderson et al., 2017). However, most research on adaptive thermal plasticity has focused on the whole-organism level, such as adult or juvenile locomotor performance, metabolic rate, behavioral thermoregulation, and thermal tolerance (Gunderson and Stillman 2015; Ruthsatz et al., 2024; Seebacher et al., 2014).

Relatively little is known about thermal plasticity in primary reproductive traits, and particularly gametes. This is an important gap, as gametes directly determine reproductive success. Sperm in particular appear to be very heat sensitive, and both spermatogenesis and the performance of sperm cells can be compromised by heat (Wang and Gunderson, 2022; Walsh et al., 2019; Snook et al., 2026). During semen production, high temperatures can cause damage across multiple sperm/ejaculate traits, including reduced motility, abnormal morphology, and lower sperm count (Sales et al., 2018, 2021; Zeh et al., 2012; Parratt et al., 2021; Chirault et al., 2015; Vasudeva et al., 2019; reviewed in Wang and Gunderson, 2022; Walsh et al., 2019; Snook et al., 2026). After production, heat exposure can decrease sperm performance and cause infertility (Chirgwin et al., 2019; Walsh et al., 2022; Rahman et al., 2009; McAfee et al., 2020; Wang et al., 2025).

Whether prolonged thermal acclimation in males triggers adaptive plastic responses in ejaculate traits is largely unknown, and existing data paint an unclear picture. Studies across vertebrates and invertebrates have found that warm acclimation induces changes in sperm morphological traits such as length (Walsh et al., 2019; Wang and Gunderson, 2022; Dougherty et al., 2024), but it is not known whether these changes are adaptive. With respect to sperm/ejaculate performance, male flour beetles (*Tribolium castaneum*) had the greatest fertility under the temperatures at which they were reared (Vasudeva et al., 2019). Conversely, male heat acclimation did not lead to improvements in sperm traits or fertilization outcomes under high temperatures in purple urchin (*Strongylocentrotus purpuratus*; Leach et al., 2021) or mosquitofish (*Gambusia holbrooki;* Adriaenssens et al., 2012). More tests of male thermal acclimation effects on sperm are clearly required before generalizations can be made.

Plastic changes in sperm could also occur via heat hardening in the post-ejaculate stage. Heat hardening is a form of rapid phenotypic plasticity in which heat tolerance increases within minutes to hours following a non-lethal heat exposure (Bowler, 2005; Loeschcke and Hoffmann, 2007; Deery et al., 2021). This response is mediated by proteins such as molecular chaperones that help protect cells from heat-induced damage (Somero et al., 2017). Heat hardening has been observed across a variety of organisms and somatic cell types (Loeschcke and Hoffmann, 2007; Deery et al., 2021; Somero et al., 2017; Angilletta, 2009) but has not been investigated for sperm of any species. It may seem that sperm heat hardening is unlikely because mature sperm are assumed to be transcriptionally quiescent (Hosken and Hodgson, 2014; Immler, 2019; Lymbery et al., 2020; Miller et al., 2005). However, recent studies have shown that semen carry stress-response proteins and untranslated mRNA for proteins including heat shock proteins (HSPs), and that sperm exhibit transcriptomic and proteomic changes in response to environmental conditions (Hosken and Hodgson, 2014; Immler, 2019; Lymbery et al., 2020, 2025; Miller et al., 2005). These findings raise the possibility for functional plasticity in sperm traits after ejaculation.

Here, we test for adaptive thermal plasticity in sperm traits based on heat exposure at both the pre-ejaculate (male thermal acclimation) and post-ejaculate (heat exposure of ejaculated sperm) stages using the brown anole lizard (*Anolis sagrei*) as a model organism. To evaluate effects of male thermal acclimation, we exposed males to warm or cool thermal regimes and repeatedly measured multiple sperm traits, including heat tolerance, over 8 weeks (Fig. 1A). We tested for a heat-hardening effect by exposing ejaculated sperm to a sublethal heat shock before measuring sperm heat tolerance (Fig. 1B). If there is adaptive thermal plasticity in sperm heat tolerance, we predicted that 1) long-term male acclimation to high temperatures would increase sperm heat tolerance, and 2) short-term sublethal heat shock would induce increased heat tolerance of ejaculated sperm.

**Fig. 1.**
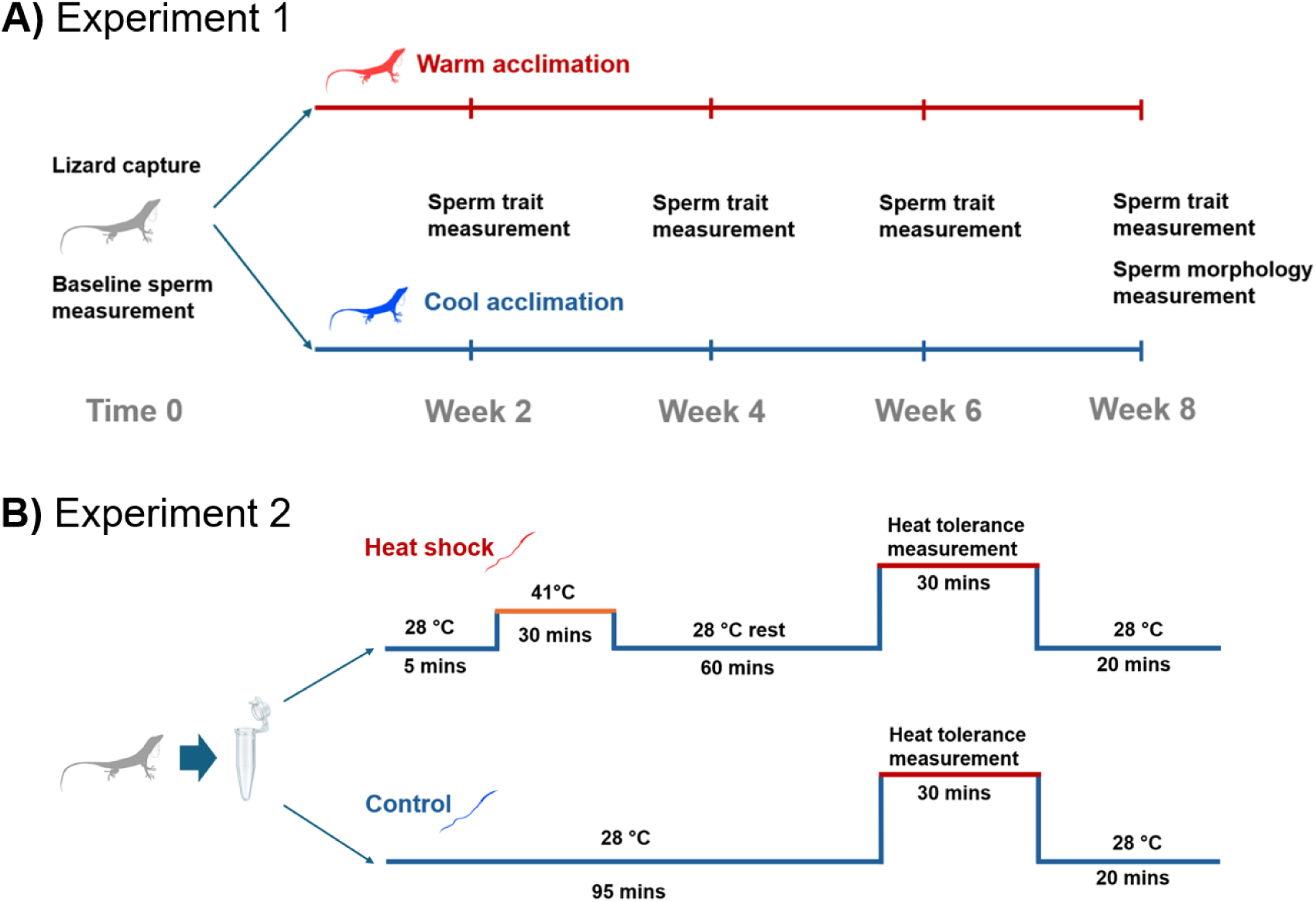
Overview of the experimental designs. (**A**) Experiment 1 was designed to test for effects of male temperature acclimation on sperm and ejaculate traits. Most traits were first measured before thermal acclimation, and then every two weeks during the eight-week acclimation period. Sperm morphology was measured at the end of the experiment (Week 8). **(B)** Experiment 2 was used to test for heat hardening in ejaculated sperm. For each male, a single ejaculate was divided into two subsamples that either did (Heat shock) or did not (Control) experience exposure to a high but non-lethal temperature (41°C) for 30 min before heat tolerance measurement.

## 2. Materials and Methods

### 2.1 Experiment 1: Testing for effects of male thermal acclimation on sperm and ejaculate traits

#### 2.1.1 Experimental Design

N = 30 adult male brown anoles were collected from the uptown area of New Orleans, Louisiana, USA (29.9407°N, 90.1203°W) in April during the breeding season. After capture, lizards were immediately brought to the lab where baseline sperm and ejaculate traits were measured (see details below). After baseline measurement, lizards were randomly assigned to one of two fluctuating thermal treatments based on ecologically relevant temperature dynamics (Figure 2) for eight weeks: "Warm" or "Cool." The "Warm" regime fluctuated between 26 and 38°C and simulated projected future conditions slightly warmer than the current mean summer body temperature dynamics of brown anoles in New Orleans (“Normal” in Fig. 2; Deery et al. 2021). The "Cool" regime fluctuated between 22-32°C and represented spring conditions early in the breeding season (Deery et al. 2021; Rej and Gunderson 2025). Lizards were housed individually in plastic cages (18×11×14 cm) in a climate-controlled growth chamber (Percival, I30NLC9) maintained on a 14:10 light:dark cycle at 75% relative humidity, watered daily and fed crickets dusted with calcium and vitamin powder three times per week. The 8-week acclimation period ensured coverage of a full spermatogenic cycle, from spermatid development to the formation of mature sperm (4∼8 weeks in reptiles; Gribbins, 2011; Rosati et al., 2022). We measured the following sperm and ejaculate traits: sperm heat tolerance, sperm motility, sperm count, and sperm morphology (see below for details). All protocols were approved by the Tulane University Institutional Animal Care and Use Committee (protocol no. 1436).

**Fig. 2.**
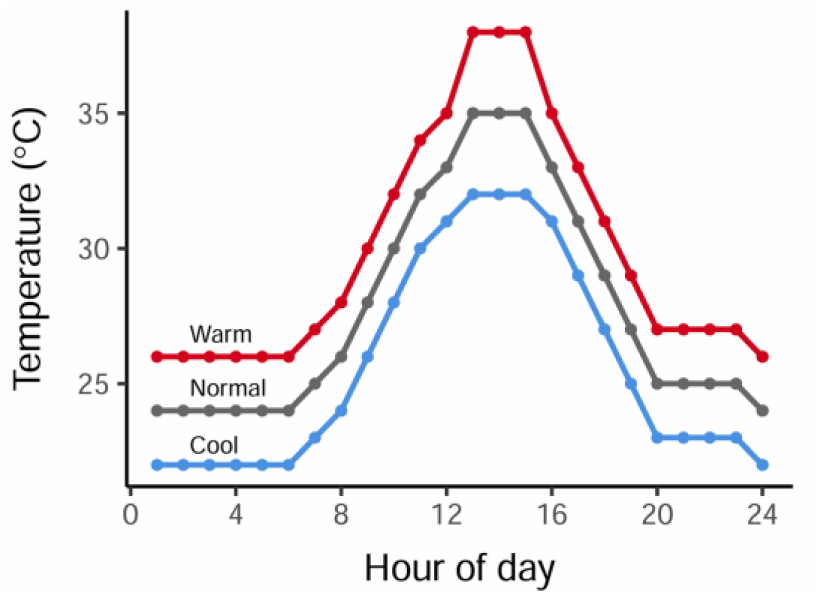
Fluctuating thermal regimes used in this study. The “Warm” acclimation regime simulated projected future temperatures above those of a typical summer day in New Orlreans (i.e., “Normal”), while the Cool acclimation regime reflected a spring day early in the reproductive season.

#### 2.1.2 Semen Collection

Semen was collected using a non-invasive technique (Martínez-Torres et al., 2019; Wang et al., 2025; Kahrl et al., 2019). Each male was gently restrained by hand, and light pressure was applied to the pelvic girdle near the cloaca until ejaculate emerged. A total of 2 μL of ejaculate was immediately collected using a pipette and diluted into 1000 μL of Dulbecco’s Modified Eagle Medium (DMEM) in a 1.5 mL Eppendorf tube. Diluted samples were then used immediately in assays.

#### 2.1.3 Sperm Heat Tolerance

Sperm heat tolerance was measured as the temperature at which 50% of sperm in an ejaculate sample become immobile (LT50) following Wang et al. (2025). The diluted semen sample from each male was divided into seven 20 μL subsamples, each loaded into a well of a different PCR plate and sealed with adhesive PCR plate seal (Thermo Scientific™). Each plate was exposed a different temperature (24°C, 37°C, 39°C, 41°C, 43°C, 45°C, or 47°C). The 24°C exposure was achieved by placing the plate on the bench in the climate-controlled lab. For the rest of the temperatures, each plate was placed inside a circulating water bath ensuring that all wells were fully submerged. Water temperature was controlled using precision immersion circulators (BLITZHOME). Multiple water baths were run in parallel, each maintained at one experimental temperature. Samples were incubated at their assigned temperatures for 30 min, after which plates were removed and allowed to cool at room temperature for 10 min. Following cooling, 10 μL from each subsample was pipetted onto a hemocytometer and immediately video-recorded for 30 sec under a compound light microscope (Nikon Eclipse) equipped with a digital camera (AmScope; MU1000-HS).

Sperm motility in each subsample was quantified using the ImageJ plugin ‘cell counter’ to calculate the proportion of sperm cells that were moving. We estimated sperm LT50 for each individual following a previously described dose–response modeling approach (Wang et al. 2025). We first normalized motility values by dividing each measurement by the maximum motility observed across all temperature treatments for that individual ejaculate. This normalization accounts for baseline variation and ensures that LT50 estimates reflect the relative decline in motility due to heat exposure rather than differences in initial motility (Ritz et al., 2015; Ritz and Streibig, 2005; Wang et al., 2025). We then fitted a four-parameter log-logistic model to the normalized motility data using the LL.4 function in the R package “drc” and LT50 was extracted from each fitted curve. To avoid bias, all steps in the experiment were randomized, including the order of sperm collection from each male, temperature assignments across PCR plates, and the order of video recording. The observer was blind to both treatment temperature and lizard identity during motility assessment.

#### 2.1.4 Sperm Count, Baseline Motility and Morphology

To determine sperm count and baseline motility, 10 μL of diluted semen was loaded into a chambered sperm counting slide (Leja slides, 20 microns, 4 chambers). A 30 sec video was recorded under 200X magnification using the same microscope and camera system described above. Motile and non-motile sperm within the field of view were manually counted using the Cell Counter plugin in ImageJ. Due to a shortage of chambered slides, baseline motility and sperm count were not measured in week 4. For sperm morphology, a separate image was taken under 400X magnification from the same slide. Morphological measurements (head, midpiece, and tail length) were taken using the software “Sperm Sizer”, and 8 to 15 individual sperm were measured per sample (McDiarmid et al., 2021; Bulla et al., 2024). Only clearly visible, non-overlapping sperm were included to ensure measurement accuracy, resulting in variation in the number of sperm measured among samples. Values for each individual sperm cell from aa given male were then averaged for subsequent analyses. Sperm morphology was measured at the end of the experiment (week 8).

### 2.2 Experiment 2: Testing for heat hardening in ejaculated sperm

N = 29 adult male brown anoles were collected from New Orleans, LA, USA during June and transported to Tulane University where they were housed under the ‘normal’ thermal regime described above (Fig. 2). After one week, semen samples were collected from each male and diluted as previously described. Diluted samples were then separated into two subsamples that each received a different thermal treatment (Fig. 1B). The Heat Shock group experienced exposure to 41°C for thirty minutes in a water bath followed by a 1h rest period at room temperature prior to heat tolerance measurement as described for Experiment 1. The temperature of 41°C was chosen for heat shock because it is the highest temperature that brown anole sperm can experience for 30 min with no effect on motility (Wang et al., 2025). The Control group only experienced 28°C, but both treatments experienced the same amount of time before and after heat tolerance measurement (Fig. 1B).

### 2.3 Replication statement

| Scale of inference | Scale at which the factor of interest is applied | Number of replicates at the appropriate scale |
| --- | --- | --- |
| Individual male (Experiment 1) | Thermal acclimation treatment (Warm vs Cool) applied to individual males. | 30 males total (Warm: 15, Cool: 15) |
| Individual male (Experiment 2) | Heat shock treatment (Heat shock vs Control) applied to split ejaculate subsamples from each male. | 29 males, each contributing one Heat shock and one Control subsample (paired within male) |

### 2.4 Data analysis

For Experiment 1, we used linear mixed-effects models (LMMs) to test for effects of male thermal acclimation treatment on sperm traits. For each trait, we calculated change in phenotype (Δ) by subtracting the baseline (Time 0) value from subsequent measurements, and used phenotype change in analyses. Each trait was modeled with thermal treatment (warm vs cool), time (week) and their interaction as fixed effects and lizard ID as a random effect to account for repeated measures. All models were fit using the “lmer” function in the “lmerTest” package in R (R Core Team, 2024; Kuznetsova et al., 2017). To test whether thermal acclimation affected sperm morphology, we compared head, midpiece, and tail length between warm and cool acclimated males using Wilcoxon rank-sum tests because data were not normally distributed (Shapiro–Wilk tests, P < 0.050).

For Experiment 2, a two-tailed paired t-test in R was used to test for differences in LT50 between Heat Shock and Control treatments after checking the normality of the data using Shapiro–Wilk tests.

## 3. Results

### 3.1 Experiment 1: Effect of male thermal acclimation on sperm and ejaculate traits

#### 3.1.1 Sperm Heat Tolerance

Sperm heat tolerance was not affected by male thermal acclimation, as sperm LT50 did not differ between males exposed to the Warm and Cool thermal regimes (Fig. 3; Table 1; p = 0.267). The mean change in LT50 from baseline to week 8 was −0.3 ± 0.5 °C in warm-acclimated males and 0.1 ± 0.4 °C in cool-acclimated males. LT50 also did not change over time (p > 0.251 for all weeks), and there were no treatment by time interactions (p > 0.768 for all weeks; Table 1).

**Fig. 3.**
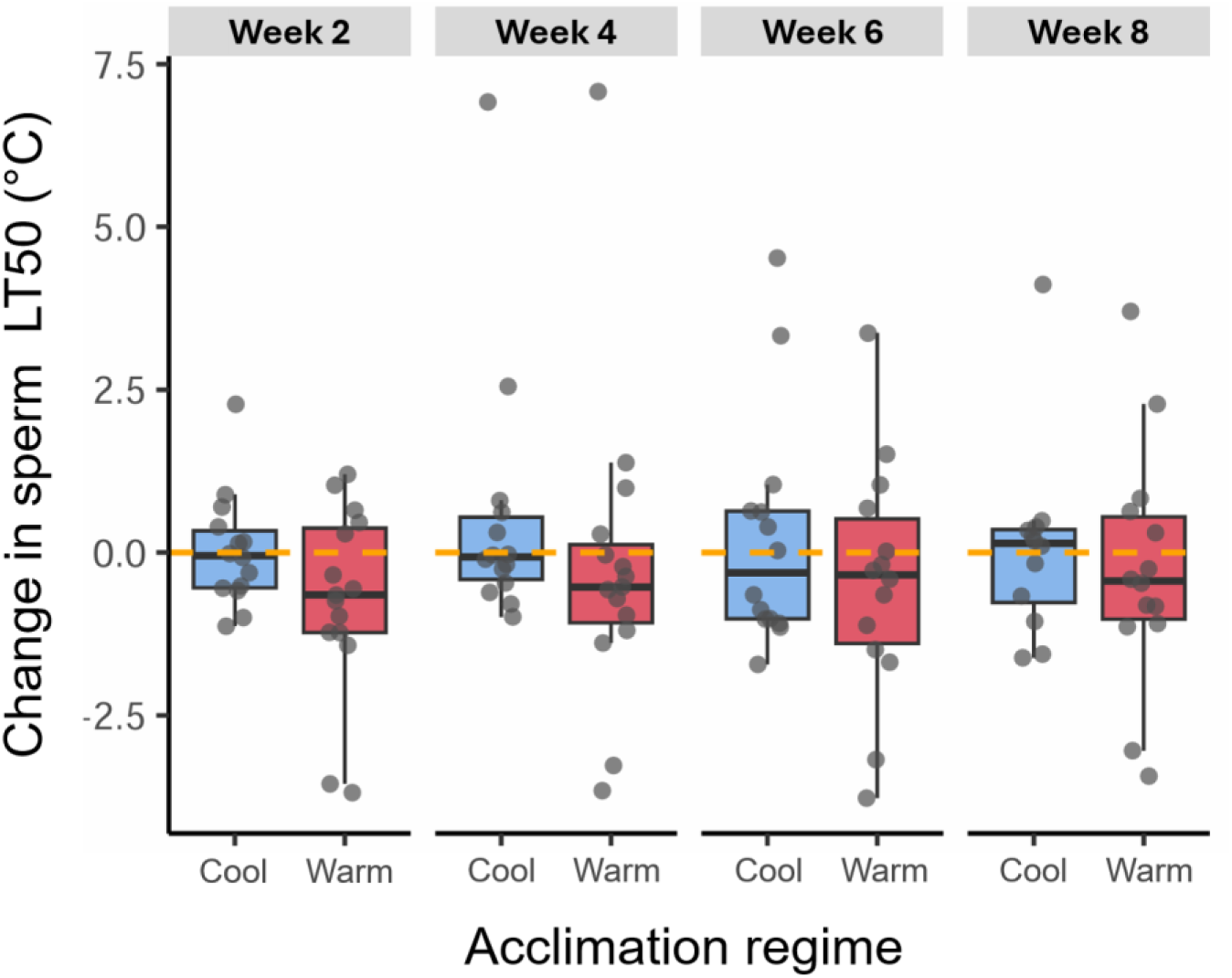
Result of male thermal acclimation on sperm heat tolerance over time. Y-axis is change in sperm LT50 relative to pre-treatment (Time 0) baseline for each individual. Each box represents the interquartile range, with the median indicated by a horizontal line. Individual data points are shown in gray. Orange dashed line indicates no change relative to pretreatment baseline.

**Table 1.** Summary of mixed-effects models testing the influence of male thermal acclimation (Experiment 1) on sperm and ejaculate traits. Each trait for an individual was centered by subtracting its baseline (Time 0) value from subsequent values. For each trait, only the smallest p-value from the group is reported. Values shown are estimated effect sizes (β ), standard error (SE), degrees of freedom (df), and *p*-values.

| | Predictor | Estimate ($\beta$) | SE | df | p-value |
| --- | --- | --- | --- | --- | --- |
| **Sperm LT50** | Treatment | -0.74 | 0.66 | 56.03 | 0.267 |
|  | Week | - | - | - | >0.251 |
|  | Week:Treatment | - | - | - | >0.768 |
| **Sperm Motility** | Treatment | +3.38 | 2.96 | 42.54 | 0.261 |
|  | Week | - | - | - | >0.583 |
|  | Week:Treatment | - | - | - | >0.524 |
| **Sperm count** | Treatment | +26.13 | 15.40 | 66.41 | 0.094 |
|  | Week | - | - | - | >0.604 |
|  | Week:Treatment | - | - | - | >0.254 |

#### 3.1.2 Sperm Motility

Sperm motility was not affected by male thermal acclimation, as it did not differ between males in the Warm and Cool treatments (Fig. 4A; p = 0.261). Mean change in sperm motility from baseline to week 8 was −1.6 ± 2.2% in warm-acclimated males and −4.4 ± 2.2% in cool-acclimated males. Motility also did not change over time (all p > 0.583), and there were no treatment by time interactions (all p > 0.524; Table 1).

**Fig. 4.**
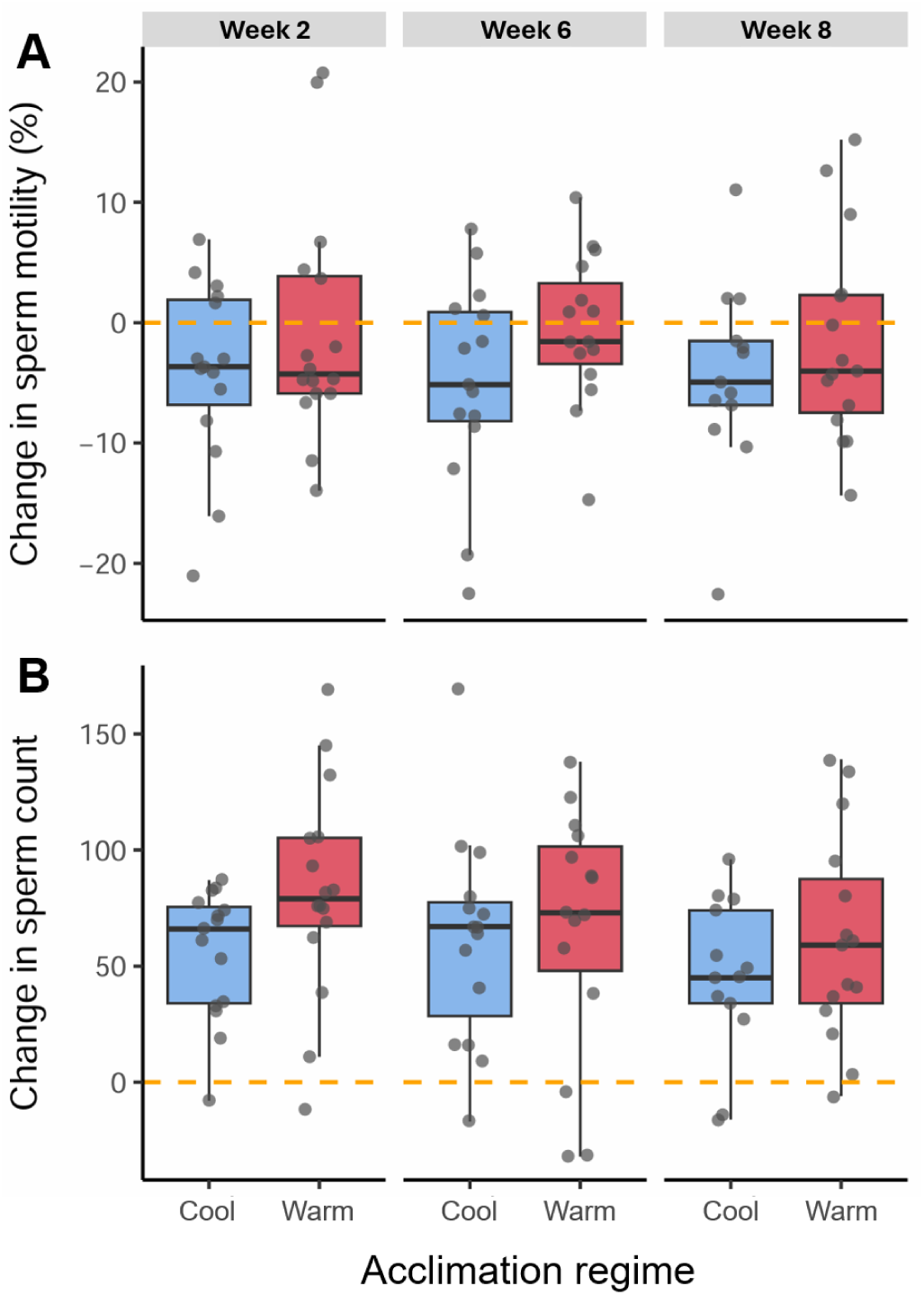
Effects of thermal acclimation on sperm motility and sperm count over time. **(A)** Change in baseline sperm motility and **(B)** change in sperm count for males acclimated to Cool (blue) or Warm (red) temperature regimes relative to their pre-acclimation baseline (Time 0). Orange dashed line indicates no change relative to baseline. Boxes show the interquartile range with medians; individual values are shown as gray points.

#### 3.1.3 Sperm Count

Sperm count generally increased across males relative to their baseline, but sperm count did not differ between males in the different thermal acclimation treatments (p = 0.094; Fig. 4B; Table 1). Mean change in sperm count from baseline to week 8 was 61.3 ± 11.6 in warm-acclimated males and 45.5 ± 9.4 in cool-acclimated males. No significant differences were observed across time (p > 0.604) and there were no treatment-by-time interactions (p > 0.254).

#### 3.1.4 Sperm Morphology

The length of the head (p = 0.323), midpiece (p = 0.303), and tail (p = 0.083) did not differ between sperm produced by males in the Warm and Cool acclimation treatments (Fig. 5).

**Fig. 5.**
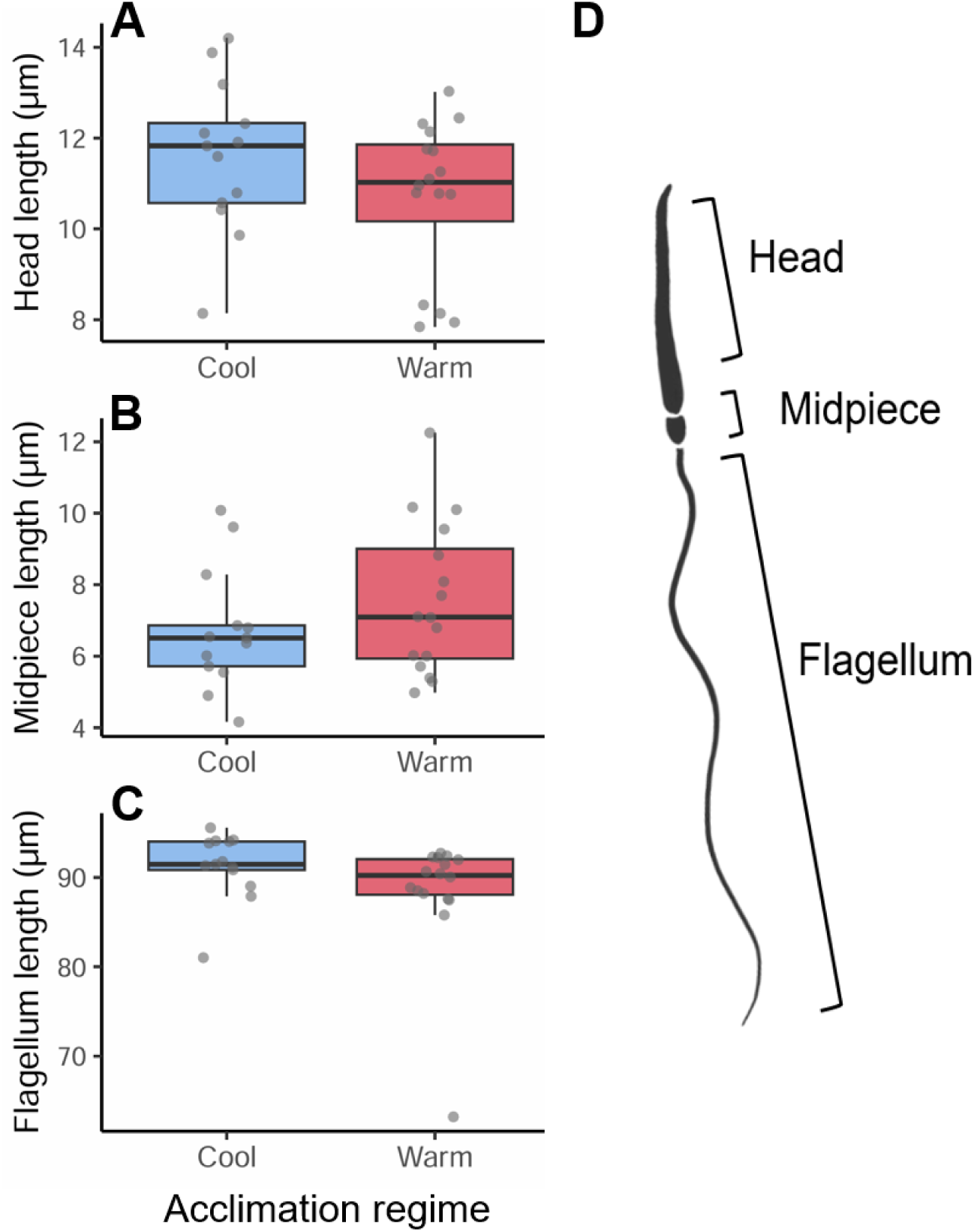
Sperm morphology by thermal acclimation treatment. **(A)** head length, **(B)** midpiece length, **(C)** flagellum length. Points represent individual lizard means. Boxes indicate interquartile ranges with medians. **(D)** Diagram of the sperm components measured. Measurements were taken at week 8.

### 3.2 Experiment 2: Effect of short-term heat shock on ejaculated sperm

Sperm LT50 did not differ between ejaculate sub-samples that were heat shocked and those that were not (Fig. 6; paired t-test: *t* = 0.30, df = 28, p = 0.77).

**Fig. 6.**
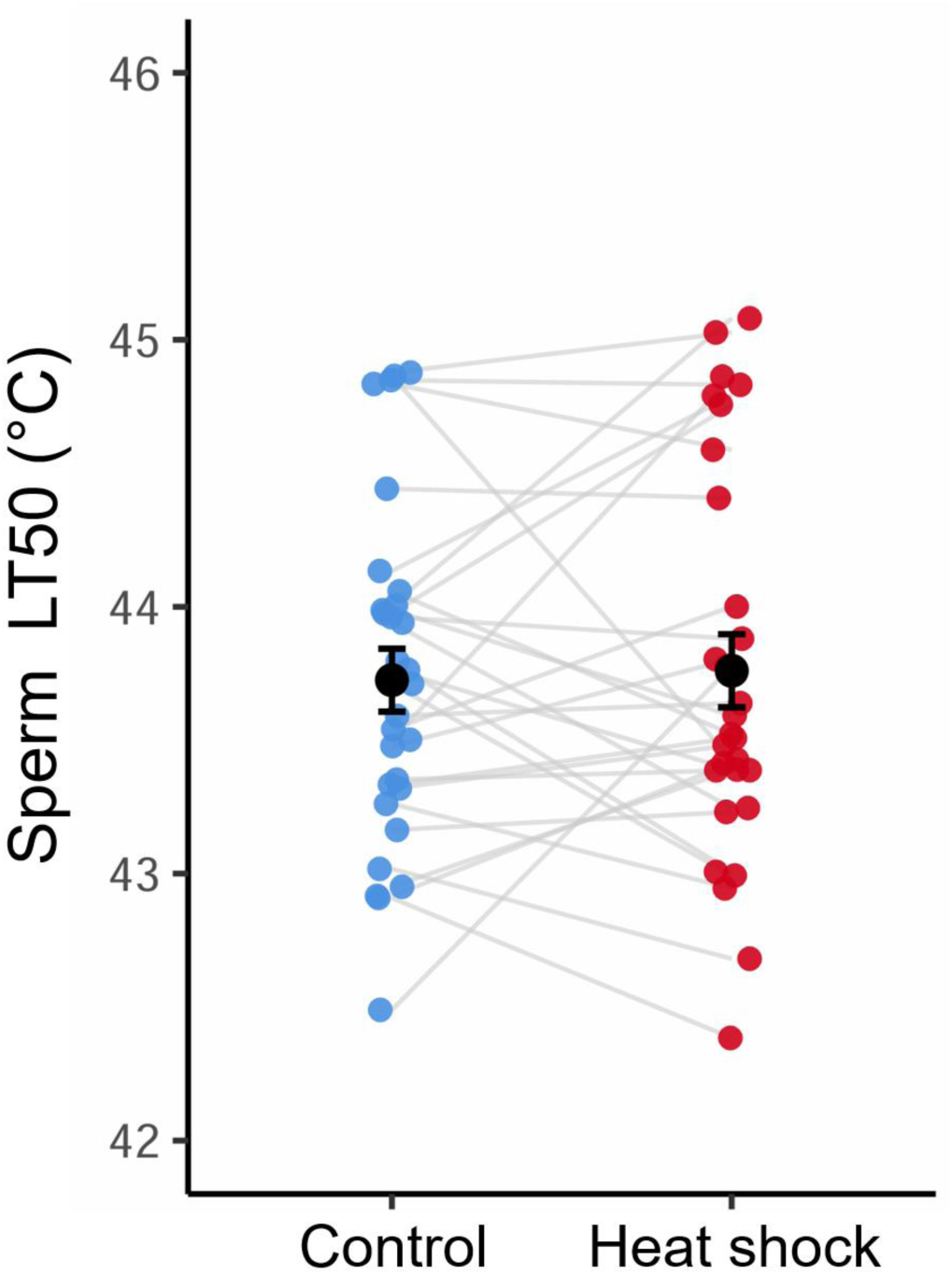
Sperm heat tolerance following heat shock. Sperm from the same ejaculate for each male were split and exposed to either a Heat shock (41°C for 30 min) or a Control condition (28°C) before heat tolerance (LT50) measurement. Black circles and bars indicate treatment means ± SE. Gray lines connect paired values from the same individual.

## 4. Discussion

Knowledge of how reproductive traits respond to thermal variation is essential for understanding thermal adaptation and predicting organismal resilience under climate change (Wang and Gunderson, 2022; Snook et al., 2026; Walsh et al., 2019; van Heerwaarden and Sgrò, 2021). However, we know relatively little about the capacity for reproductive traits to exhibit adaptive thermal plasticity, and especially at the gamete level, because most of the work on thermal plasticity has focused on adult whole-organism traits. Here, we tested for thermal plasticity in sperm and ejaculate traits in response to male thermal acclimation (Experiment 1), and tested whether exposing ejaculated sperm cells to heat induces heat hardening (Experiment 2). We discuss these results in turn below.

Contrary to predictions from adaptive plasticity, we found no evidence that male acclimation to heat improves the heat resilience of brown anole sperm. Males exposed to a high and ecologically realistic fluctuating temperature regime for eight weeks did not produce sperm cells with greater heat tolerance (Fig. 3; Table 1). Experimental tests for effects of male thermal acclimation on sperm cell thermal performance are rare (reviewed in Wang and Gunderson, 2022; Walsh et al., 2019). However, our findings align with several previous studies in other taxa, including in mosquitofish (*Gambusia holbrooki;* Adriaenssens et al., 2012; Iglesias-Carrasco et al., 2020), brown trout (*Salmo trutta;* Fenkes et al., 2017) and European whitefish (*Coregonus lavaretus*; Kekäläinen et al., 2018), which found that male thermal environment had limited or no effect on sperm performance under thermal stress, with no evidence that acclimation enhances sperm heat tolerance.

We are aware of only one study that has demonstrated adaptive thermal plasticity in male sperm performance, wherein male flour beetles (*T. castaneum*) reared at warm temperatures exhibited higher fertility under warm conditions (Vasudeva et al., 2019). Importantly, the male beetles were reared under their respective thermal treatments from hatching, meaning effects of developmental plasticity and reversible adult acclimation cannot be teased apart (Vasudeva et al., 2019). We tested for reversible acclimation over eight weeks during adulthood, a duration that is sufficient to encompass a full spermatogenic cycle in vertebrates (Sharma and Agarwal, 2011; Gribbins, 2011; Rosati et al., 2022). Further studies are needed to test whether temperature exposure during early life is more likely to induce adaptive plastic responses in sperm thermal traits across a wide range of organisms.

Consistent with our sperm heat tolerance results, none of the other ejaculate and sperm traits that we measured were affected by male thermal acclimation, including sperm motility, count, and morphology (Fig. 4, 5; Table 1). These results are somewhat surprising, as other studies have reported changes in these types of traits following male exposure to elevated temperatures (reviewed in Wang and Gunderson 2022, Walsh et al., 2019). For example, in the southern bunchgrass lizard (*Sceloporus aeneus*), exposing males to high temperatures for just a week increased sperm morphological abnormalities, decreased sperm viability, lowered sperm concentration, and reduced motility (Quintero-Pérez et al., 2023). Several studies have also found that males exposed to or reared at high temperatures produce shorter sperm (Wang and Gunderson, 2022; Vasudeva et al., 2019; Breckels and Neff, 2013; Adriaenssens et al., 2012; Iossa), though whether sperm morphological changes in response to temperature are adaptive remains largely unknown given a lack of direct tests linking sperm length to fertilization success across different thermal conditions.

A key difference between our approach and that of most previous studies of thermal plasticity in sperm traits is the dynamic temperature regimes that we used. The males in our thermal acclimation treatments experienced daily temperature fluctuations that reflect what these animals experience in the wild. In contrast, most previous studies acclimated animals to constant temperatures. It is well established that the consequences of temperature exposure, including physiological plasticity, can differ when animals are experience fluctuating versus constant temperatures, even when mean temperatures are the same (Niehaus et al., 2011; Marshall et al., 2021). The difference in approach could partially explain why we do not see changes in sperm and ejaculate traits that others have found. We recommend that future studies employ more ecologically relevant experimental regimes that reflect the dynamic conditions that animals experience in the wild to assess reproductive performance and plasticity in response to heat (Wang and Gunderson, 2022).

Rapid heat hardening is a common form of putatively adaptive thermal plasticity common at both the whole-organism and cellular level (Loeschcke and Hoffmann, 2007; Deery et al., 2021; Somero et al., 2017; Angilletta, 2009). Sperm are thought to be transcriptionally silent due to chromatin condensation and histone replacement during spermatogenesis (Hosken and Hodgson, 2014; Miller et al., 2005; but see Lymberry et al. 2025, 2020), but this does not necessarily preclude rapid physiological changes. Sperm contain heat shock protein mRNAs, heat exposure changes the mRNA profile of sperm cells (Lymbery et al., 2020), and many of the cellular changes that accompany rapid heat hardening result from the activity of enzymes that are present in cells and do not require transcriptional change (Hochachka, 2019; Somero et al., 2017). Despite this potential, we found no evidence of heat hardening in ejaculated sperm (Fig. 6). To our knowledge, our study is the first to explicitly test for rapid heat hardening in sperm cells, but our finding is consistent with previous studies testing for adaptive thermal plasticity in sperm cells after ejaculation. For example, in European whitefish (*C. lavaretus*) and mussels (*M. galloprovincialis*), exposing ejaculated sperm cells to warmer activation temperature or pre-exposing them to warmer temperatures did not induce plastic improvements in their motility in warmer water (Kekäläinen et al., 2018; Lymbery et al., 2020). Our results indicate that the ability of sperm to upregulate stress responsive processes under heat shock is constrained (Lymbery et al., 2020).

Our finding that sperm traits lack adaptive plasticity has important implications for species’ persistence under climate change. Because reproductive success can be directly constrained by sperm function (Gomendio et al., 2007; Gage et al., 2004; Sales et al., 2018, 2021), the absence of within-individual plasticity means that males may not be able to buffer fertilization performance against acute or chronic warming. It appears that ectotherm sperm traits have less thermal plasticity than whole-organism thermal tolerance traits that are themselves known to have limited capacity for adaptive heat tolerance plasticity (Gunderson and Stillman, 2015; Seebacher et al. 2014; Pottier et al. 2022, 2025; Ruthsatz et al. 2024). If the thermal plasticity of reproductive traits is low, other mechanisms would be required to buffer reproduction against ongoing environmental warming, such as behavioral heat avoidance and evolutionary change. Information on the evolvability of sperm heat tolerance is extremely limited, but previous work has found moderate repeatability in the heat tolerance of brown anole sperm cells, indicating potential for evolutionary change (Wang et al., 2025).

It is important to note that our results do not preclude other forms of adaptive thermal plasticity in sperm performance. As mentioned above, adaptive developmental plasticity is a possibility (Vasudeva et al., 2019). In addition, sperm performance in internal fertilizers can be shaped by the female reproductive environment, including interactions between seminal fluid proteins, ovarian fluid and other chemical factors within the female reproductive tract (Wolfner, 2011; Pitnick et al., 2020; Holt and Fazeli, 2016; Rossi et al., 2021; Kustra et al., 2026). Therefore, it is possible that thermal acclimation modifies ejaculate–female interactions in ways that enhance sperm performance under thermal challenges. Females can also adjust their mating behaviors to compensate for lower male fertility under heat (Vasudeva et al., 2021; Pembury Smith et al., 2025). Future work integrating ejaculate–female interactions will be important for understanding the full scope of thermal plasticity in reproductive performance.

## 5. Conclusion

We conducted a robust study to directly test for adaptive thermal plasticity in sperm cells and ejaculate traits under ecologically relevant fluctuating thermal conditions. We found no evidence for adaptive thermal acclimation in the primary male reproductive traits of brown anoles. We also found no evidence of heat hardening in ejaculated sperm. This suggests that gamete-level traits have limited capacity for within-individual adjustment to thermal stress, potentially increasing reliance on evolutionary and behavioral responses to climate change. Nonetheless, this remains a largely unexplored area that warrants further investigation.

## Acknowledgements

We thank all members of the Gunderson Lab for their assistance and for valuable discussions throughout this project. We are especially grateful to Rou-Rung Chen, Natalie R. Page, Anthony M. Strickler, Jalen LaCour, and Jess Valentine. We thank Dr. Scott Pitnick, Dr. Kathleen Ferris, Dr. Jordan Karubian, Dr. Ariel Kahrl for their guidance, constructive feedback, and insightful discussions. We also thank the Department of Ecology and Evolutionary Biology at Tulane University for funding and institutional support, and are grateful to all colleagues and collaborators for helpful comments and conversations during the development of this work.

## Author contributions

Wayne Wen-Yeu Wang and Alex R. Gunderson conceived the ideas and designed methodology; Wayne Wen-Yeu Wang, Benjamin Pethe, Waverly Wood, Colin T. Smith, Alanna J. Frick, Shannan S. Yates and Abigail Ward collected the data; Wayne Wen-Yeu Wang analyzed the data; Wayne Wen-Yeu Wang and Alex R. Gunderson led the writing of the manuscript. All authors contributed critically to the drafts and gave final approval for publication.

## 7. Supplementary

**Fig. S1.**
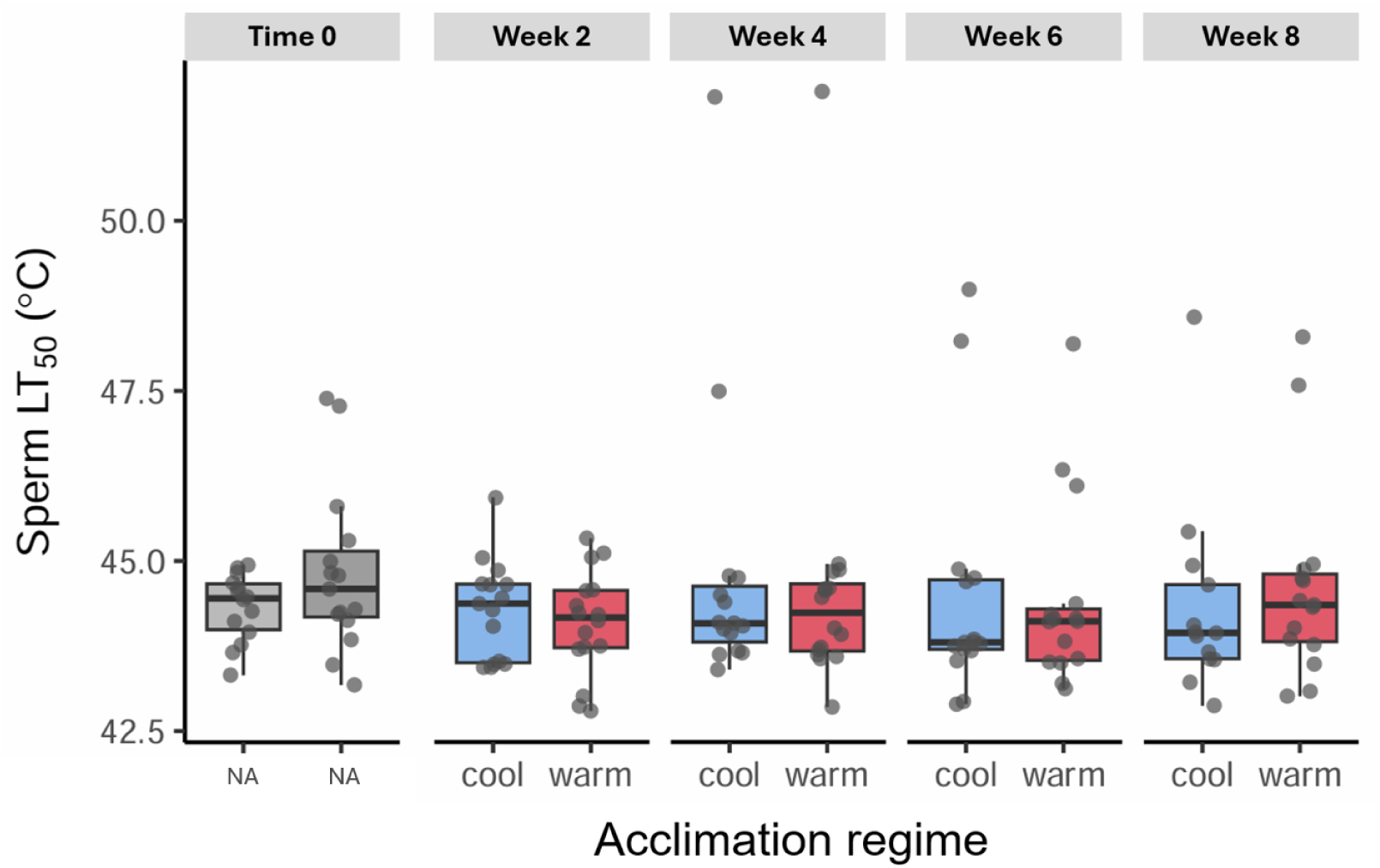
Raw sperm heat tolerance (LT50) across thermal acclimation treatments and time. Sperm heat tolerance (LT50) for males acclimated to cool and warm thermal regimes across the experimental period. Values shown here are raw LT50 measurements rather than change relative to baseline (Δ). Time 0 (gray) represents baseline values before treatment. Boxes show interquartile ranges with medians; individual values are shown as gray points.

**Fig. S2.**
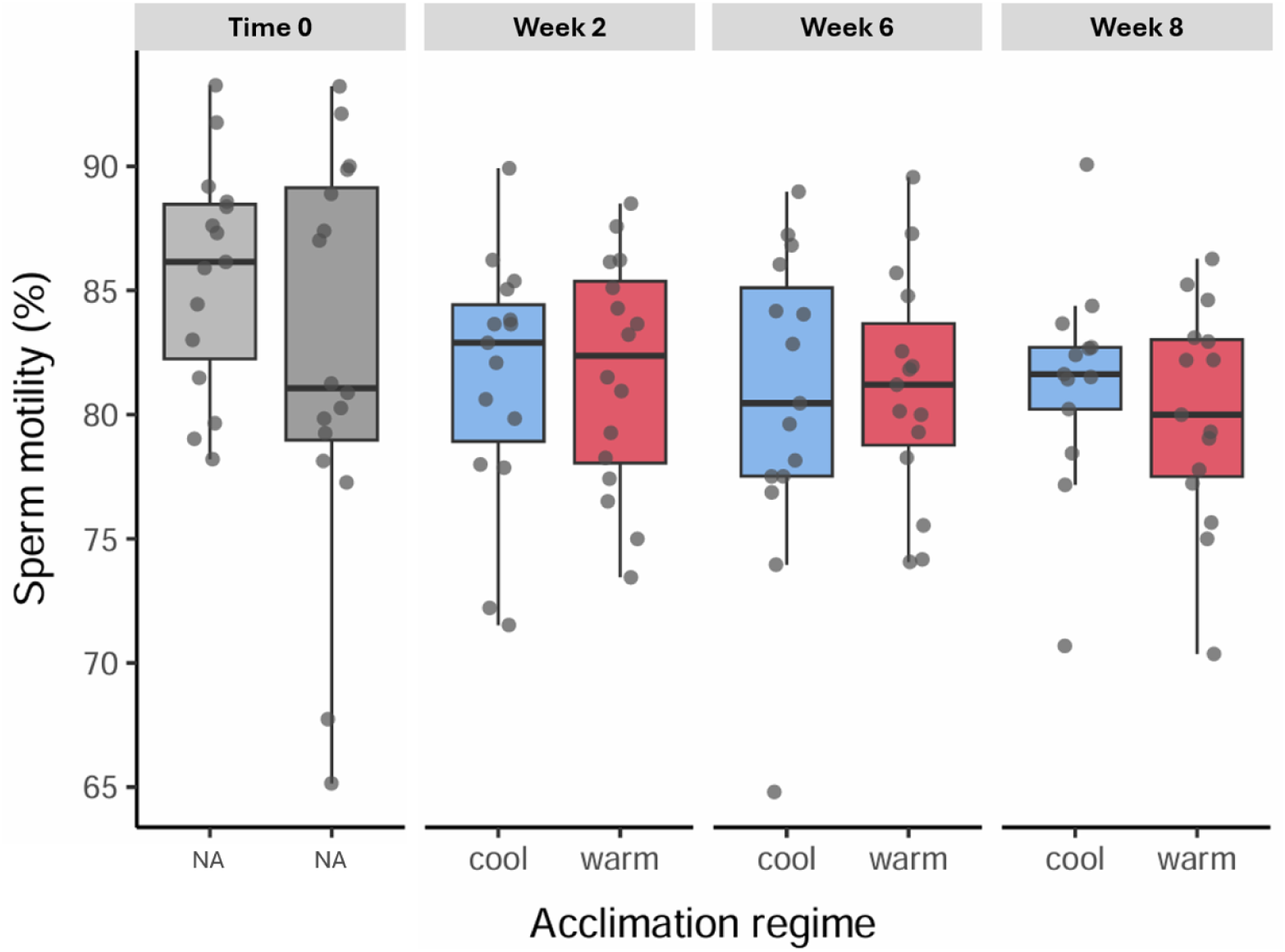
Raw baseline sperm motility across thermal acclimation treatments and time. Sperm motility (%) for males in the cool and warm acclimation treatments across time. Values are presented as raw measurements rather than Δ from baseline. Time 0 (gray) indicates pre-treatment values. Boxes represent interquartile ranges with medians; individual values are shown as gray points.

**Fig. S3.**
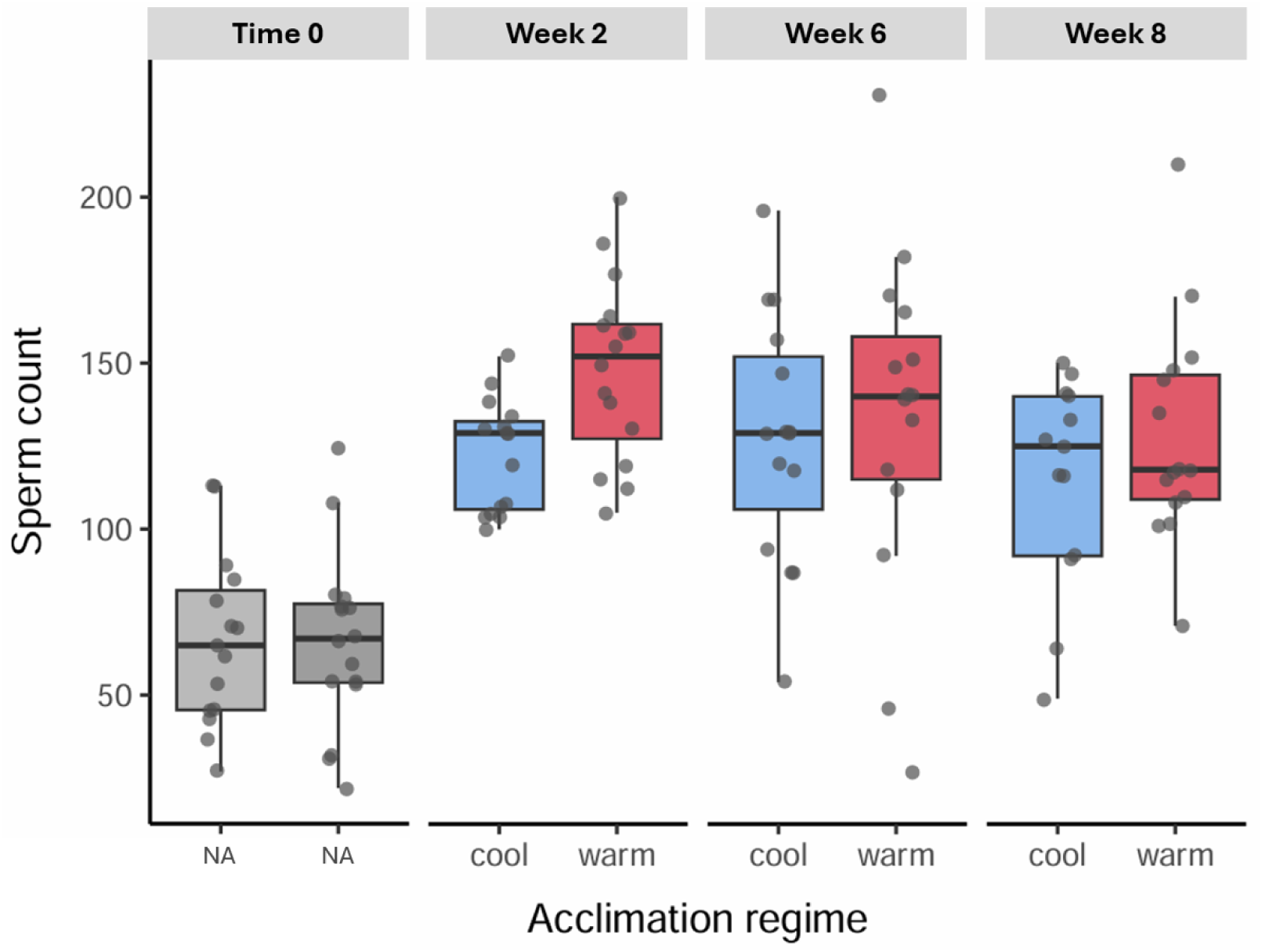
Raw sperm count across thermal acclimation treatments and time. Sperm count for males in the cool and warm acclimation treatments measured across the experiment. Values shown here are raw sperm counts rather than change from baseline (Δ). Time 0 values (gray) were collected prior to treatment. Boxes show interquartile ranges with medians; individual values are shown as gray points.

